# Torpor energetics are related to the interaction between body mass and climate in bats of the family Vespertilionidae

**DOI:** 10.1101/2023.09.30.560312

**Authors:** Jorge Ayala-Berdon, Kevin I. Medina-Bello

## Abstract

Torpor is an adaptive strategy that allows animals to cope with energy limitations under adverse environmental conditions. In birds and mammals, intrinsic and extrinsic factors such as body mass (*M_b_*) and ambient temperature (*T_a_*) are well established triggers of torpor. Interestingly, the interplay between *M_b_* and climate with different *T_a_* on torpor traits in bats remains unexplored. Using open flow respirometry, we calculated *T_a_* upon entering torpor (*T_a_t*), the reduction in torpid metabolic rate relative to the basal metabolic rate (*TMR_red_*), the *T_a_* at which torpor metabolic rate reached its minimum (*T_a adjust_*), and minimum torpid metabolic rate (*TMR_min_*) in 11 bat species of the family Vespertilionidae that differ in *M_b_* from warm and cold climates. We also included *TMR_min_* data retrieved through a bibliography review. We tested the effects of *M_b_* and climate on torpor traits using mixed-effect phylogenetic models. All models showed a significant interaction between *M_b_* and climate. This interaction was inversely related to *T_a_t*, *TMR_red_*, *T_a adjust_*, and positively related to *TMR_min_*. These results are likely explained by the differences in *M_b_* and the metabolic rate of bats from different climates, which may allow individuals to express torpor in places with different *T_a_*. Further studies to assess torpor use in bats of different climates are proposed.

**Summary statement:** The interaction between body mass and climate influences torpor energetics in bats of the family Vespertilionidae. As a result, torpid traits change based on body mass and climate.

## Introduction

Torpor is an adaptive strategy that enables animals to cope with energy limitations under adverse environmental conditions (Geiser, 2004; Geiser and Baudinette, 1988; Geiser and Brigham, 2012; Heldmaier et al., 2004; Körtner and Geiser, 2000). This physiological response is characterized by a controlled and reversible depression of animals’ metabolic rate, accompanied by a decrease in their body temperature (*T_b_*) (Fig. 1) (Geiser, 1988; Geiser, 2004; Geiser and Ruf, 1995). By using torpor, individuals can reduce their energy expenditures by 50–90 % while passively rewarming to return to normothermia (i.e., when utilizing external heat sources such as basking in the sun) (Geiser, 2008; Warnecke et al., 2008; Körtner and Geiser, 2009; Pretzlaff et al., 2010). It is well documented that intrinsic factors such as body mass (*M_b_*), as well as extrinsic factors such as ambient temperature (*T_a_*) and food availability are the primary selection pressures that trigger torpor in birds and mammals (McNab, 1989; Geiser and Brigham, 2000; Willis et al., 2005; Machado and Soriano, 2007; Wojciechowski et al., 2007; Doucette et al., 2012). Nevertheless, torpor can also facilitate energy balance during high energy-demand periods such as migration (Carpenter and Hixon, 1988; McGuire et al., 2014), development (Geiser, 2008), pregnancy (Willis et al., 2006), lactation (Chruszcz and Barclay, 2002), environmental disasters (Stawski et al., 2015) or situations involving high predation risk (Stawski and Geiser, 2010).

**Figure 1.**
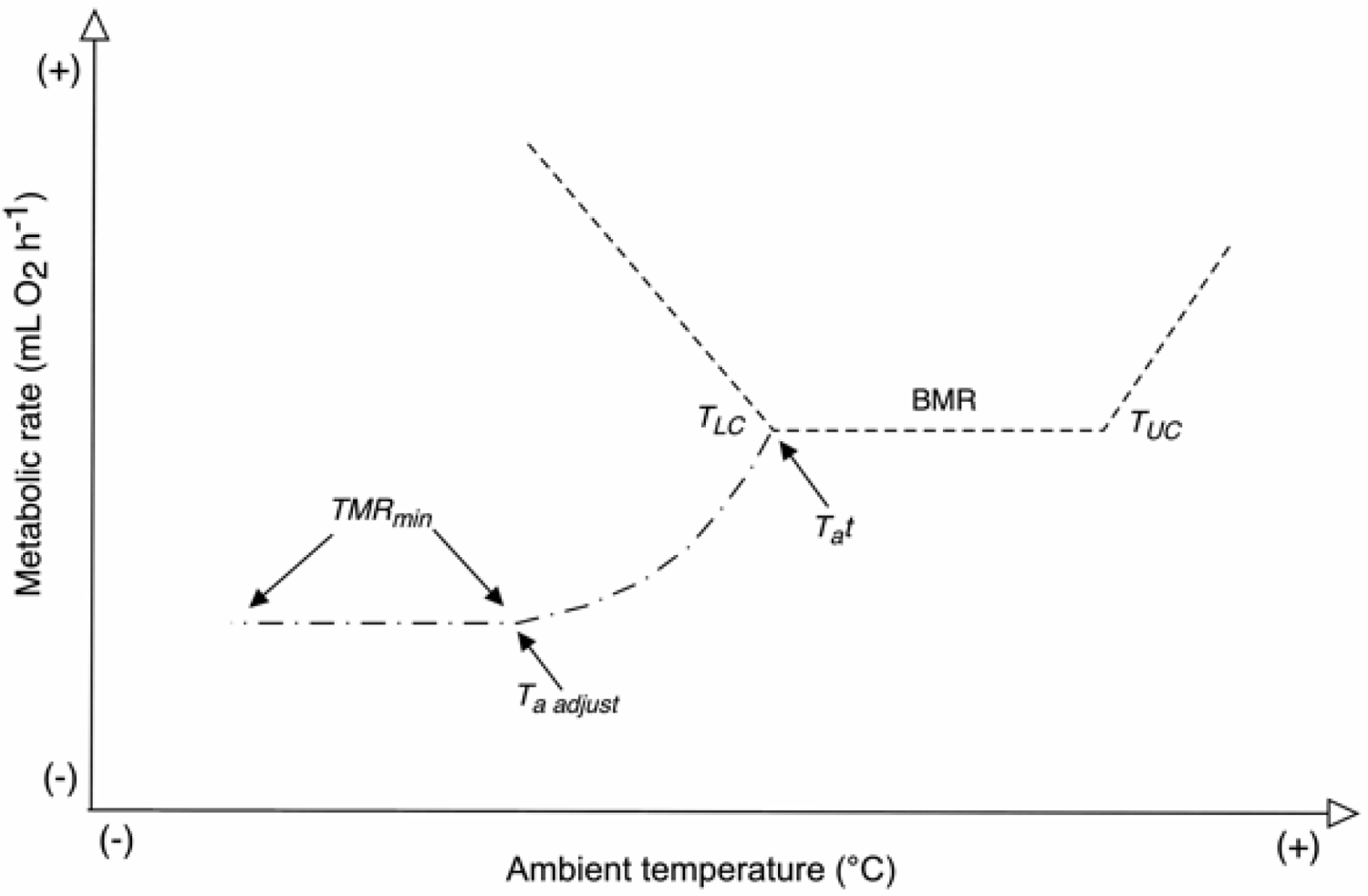
Conceptual model proposed by Speakman and Thomas (2003) of torpor energetics in heterothermic animals. In this representation the first inflection point of the metabolic rate below lower critical temperature (*T_LC_*) indicates the *T_a_* at which animals enter torpor (*T_a_t*). As *T_a_* continues to drop, the metabolic rate reaches a second inflection point (*T_a adjust_*), where the torpid metabolic rate starts reaching its minimum values (*TMR_min_*).

**Figure 2.**
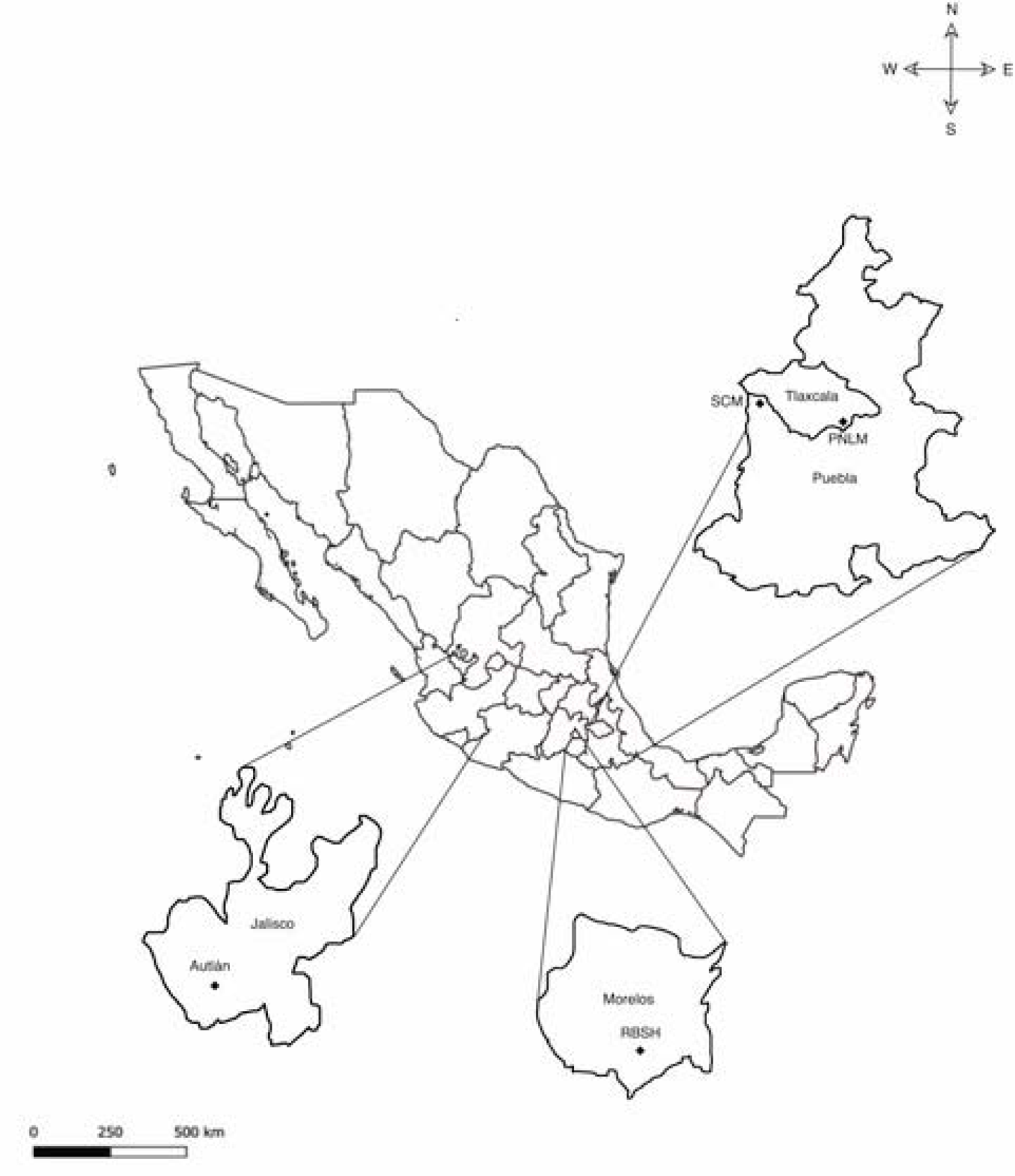
Bats were captured in four locations with either cold or warm climates in central Mexico; La Malinche National Park (LMNP) and Santa Cruz Moxolahuac (SCM) were the cold sites and Sierra de Huautla Biosphere Reserve (SHBR), and Autlán were the warm sites.

Torpor can be measured under controlled conditions (Geiser, 2021). Laboratory experiments have shown that in post-absorptive euthermic adult non-reproductive animals, a typical response to a decrease in *T_a_* below the lower critical temperature (*T_LC_*) is to increase resting metabolic rate to prevent hypothermia (Young et al., 1989). However, when environmental conditions are severe, animals may use torpor, thus reducing their metabolism below the basal metabolic rate (*BMR*) (Song et al., 1995; Geiser, 2004). Basal metabolic rate represents the minimal metabolic rate to sustain thermoregulation during normothermia and is limited by the *T_LC_* and the upper critical temperature (*T_UC_*) (representing the *T_a_* where animals begin expending energy to avoid hyperthermia) (Fig. 1) (Terrien et al., 2011). Speakman and Thomas (2003) proposed that the relationship between the metabolic rate and *T_a_* can be similar during torpor and normothermia. For instance, when individuals experience *T_a_* below their *T_LC_*, they seem to abandon normothermia by regulating *T_b_* at a very low rate, as if they were thermoconforming. In this scenario, the first inflection point of the torpor metabolic rate that falls below the *BMR* indicates the *T_a_* at which animals enter torpor (*T_a_t*). As *T_a_* continues to drop, the metabolic rate reaches a second inflection point (*T_a adjust_*), where torpid metabolic rate approaches its minimum values (*TMR_min_*). Speakman and Thomas (2003) found a positive relationship between residual *TMR_min_* (the metabolic rate that was not explained by *T_b_*) and body mass (*M_b_*) in 18 species from five families of bats. The authors also demonstrated that the minimum *T_b_* of animals during torpor (*T_bmin_*) was higher in species inhabiting warmer climates compared to colder ones. Interestingly, *T_bmin_* has also been positively related to *M_b_*in other species of birds and mammals (Ruf and Geiser, 2015; Geiser, 2021). Given that *T_b_* is closely related to the metabolic rate of animals, this strongly suggests that *TMR_min_* and other torpor traits, such as those described above, might vary among bats with different *M_b_* inhabiting warm and cold climates. Nevertheless, to the best of our knowledge, no empirical attempts have been made to assess the effect of the relationship between *M_b_* and climate on torpor traits in any mammal. This is surprising, as torpor energetics significantly influence many other aspects of species’ biology, ecology, and geographical distribution (Law, 1994).

Bats are an especially well-suited group in which to explore the effects of *M_b_* and climate on torpor energetics. They exhibit an exceptionally wide range of *M_b_* among species—from the ∼1.5 g bumblebee bat (*Craseonycteris thonglongyai*) to the ∼1,500 g giant golden-crowned flying fox (*Acerodon jubatus*) (Stawski et al., 2014). Furthermore, bats are the mammal group with the highest proportion of species that employ torpor as an energy-saving strategy across many environmental conditions (Stawski et al., 2014; Czenze and Dunbar, 2020; Geiser, 2021;). Within this group, bats from the family Vespertilionidae: i) differ in *M_b_*by approximately one order of magnitude (Moratelli et al., 2019), ii) have distributions spanning both cold and warm environments (Nowak and Walker, 1994), and iii) most species, regularly use torpor in a variety of conditions (Audet and Fenton, 1988; McWilliam, 1988; Geiser and Brigham, 2000; Genoud and Christe, 2011). In vespertilionid bats, torpor traits could be influenced by *M_b_*, climate, and/or the interaction among these explanatory variables. To test this hypothesis, we evaluated the relationship among torpor traits, *M_b_*, and climate in bats of the family Vespertilionidae that differ in *M_b_* from warm and cold climates using comparative phylogenetic models. We predicted a positive relationship between *M_b_*and torpor traits (*T_a_t*, *T_a adjust_* and *TMR_min_*). Nevertheless, bats from the warmer sites should present higher values of torpor traits compared to bats from the colder ones.

## Material and methods

### Study area

Bats were captured in four locations with either cold or warm climates in central Mexico (Fig. 1): La Malinche National Park, in Tlaxcala state, and Santa Cruz Moxolahuac, in Puebla state (colder sites); and Sierra de Huautla Biosphere Reserve, in Morelos state, and Autlán, in Jalisco state (warmer sites). Climate classification followed Köppen’s categorization, considering both the *T_a_* and precipitation of the capture sites. In this classification, warm climates exhibit a mean annual *T_a_* between ∼ 24 to 40 °C, with the minimum annual *T_a_* above10 °C, and annual precipitation ranging between 600- and 2,000-mm. Cold climates have an average annual *T_a_* between ∼ 10 and 22 °C, an annual minimum *T_a_* that tends to fall below ∼ 0 °C, and annual precipitation that ranges between ∼ 400 and 1,400 mm (Beck et al., 2005; Oliver, 2008) (Table 1).

**Table 1.**
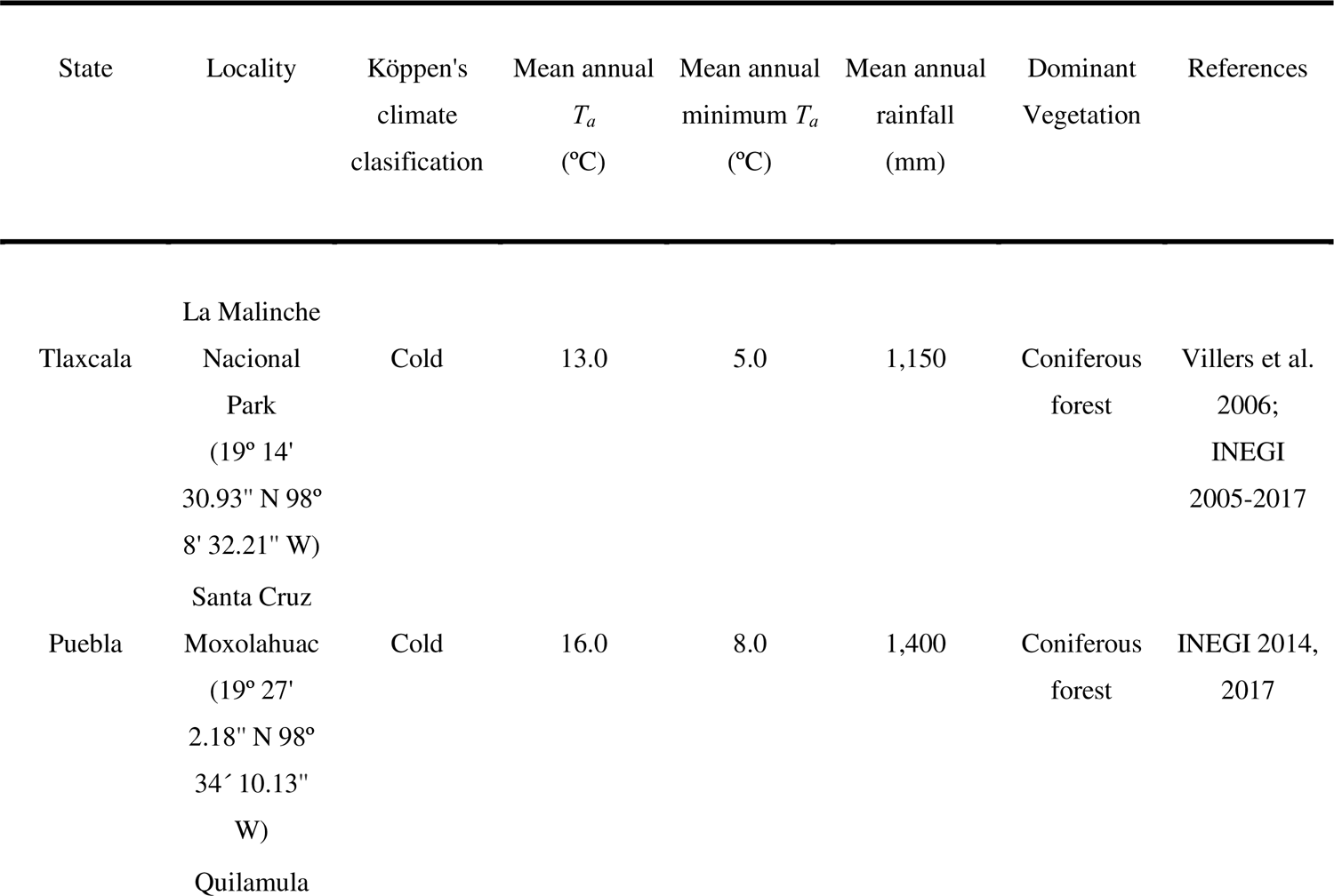
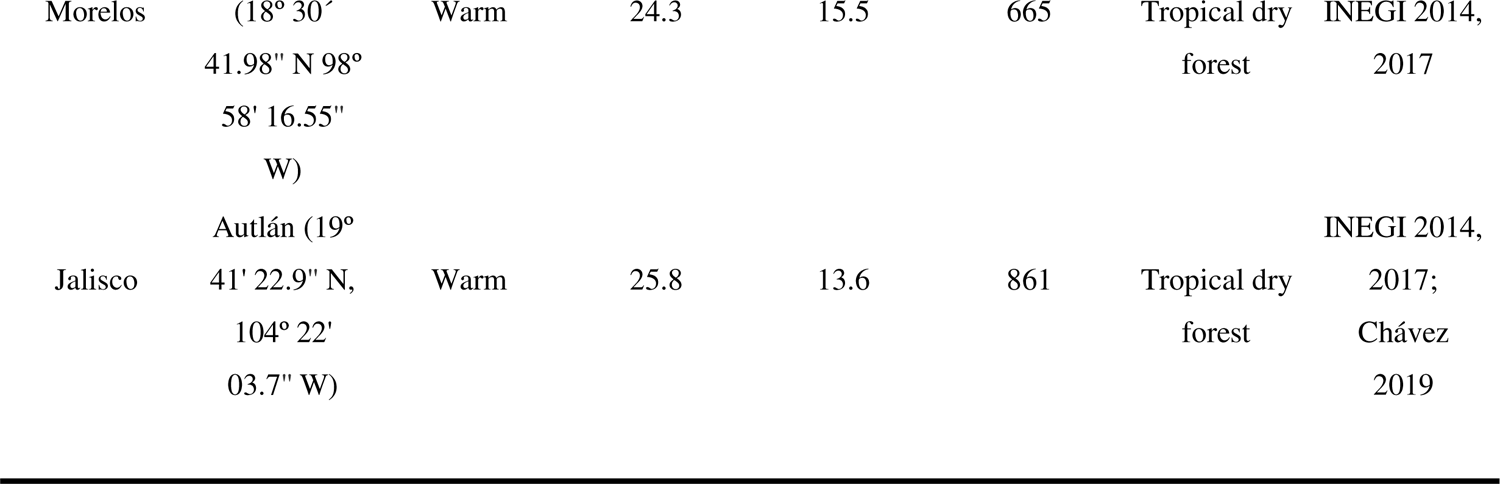
Environmental variables of the capture sites for the measurements of torpor energetics of bats of the family Vespertilionidae from central Mexico. Climate classification followed Köppen’s categorization, considering both *T_a_* and precipitation.

### Bat care and housing

Bats were captured between September and February 2021 and 2022 using mist nets (2 x 3 m, 2 x 6 m and 2 x 9 m) deployed for two consecutive nights each month at each locality during the new moon period near water bodies used by bats to drink and feed. Nets were opened at dusk and closed at ∼ 01:00 am. Adult non-reproductive males of 11 bat species of the family Vespertilionidae that vary in *M_b_*, were selected (Table 2). Some species were caught in limited numbers due to their low population size at the study localities (LaVal, 2004; Perry and Carter, 2010). Species identification followed Medellín et al. (2008). Taxonomic names were assigned following Ramírez-Pulido et al. (2014). We recorded the *M_b_* of each bat to the nearest 0.2 g using an electronic balance (Ohaus, Newark, NJ, USA).

**Table 2.**
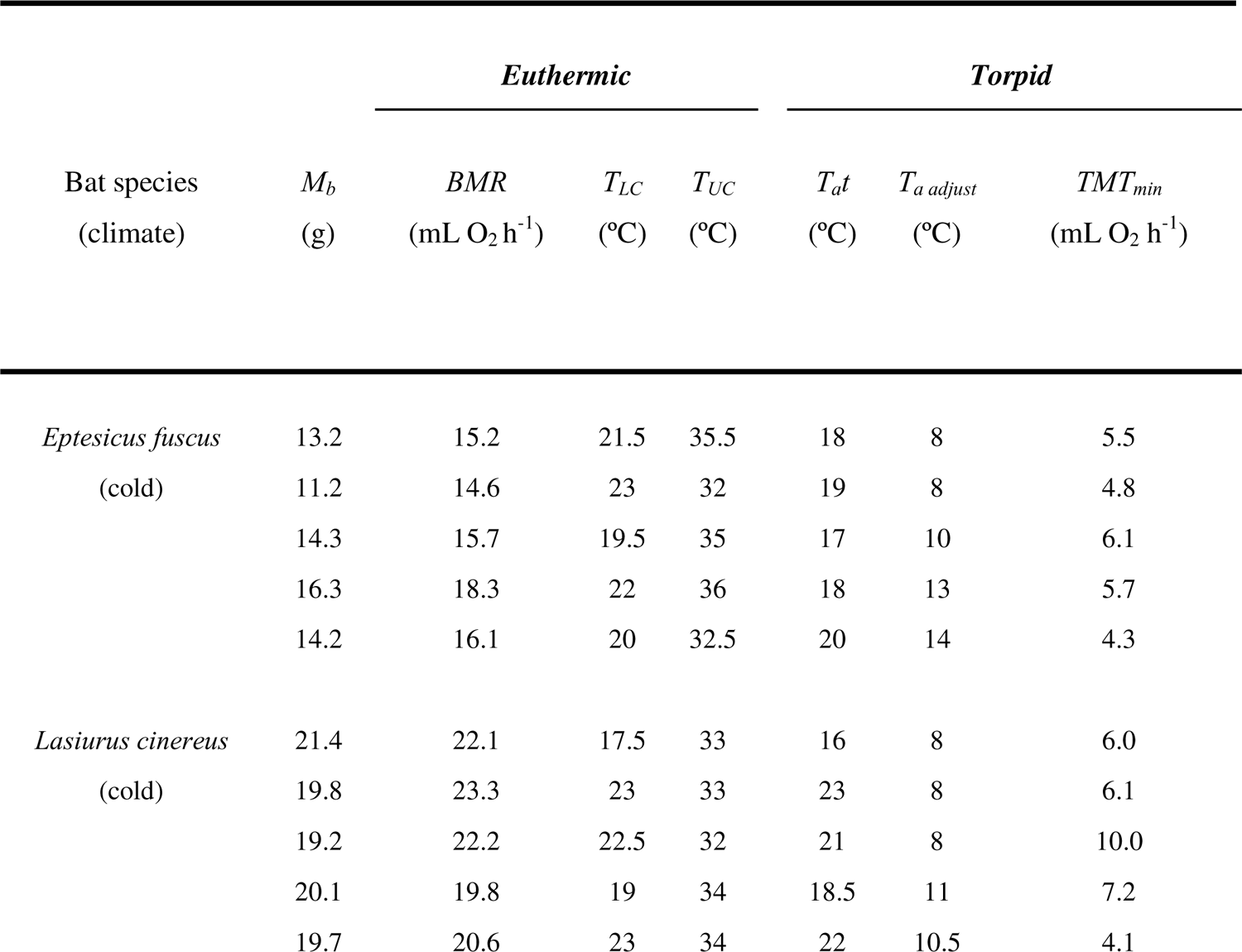
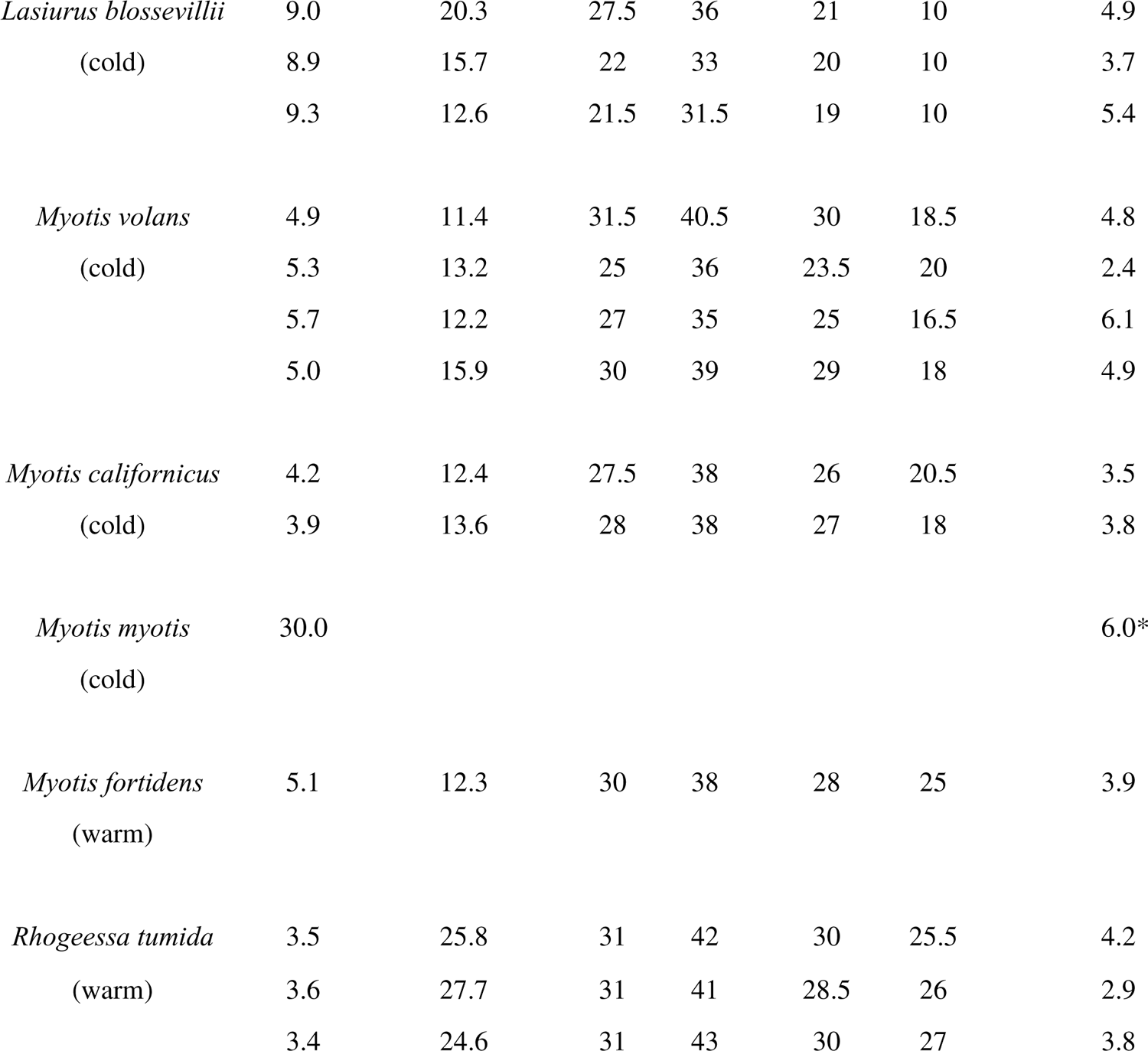
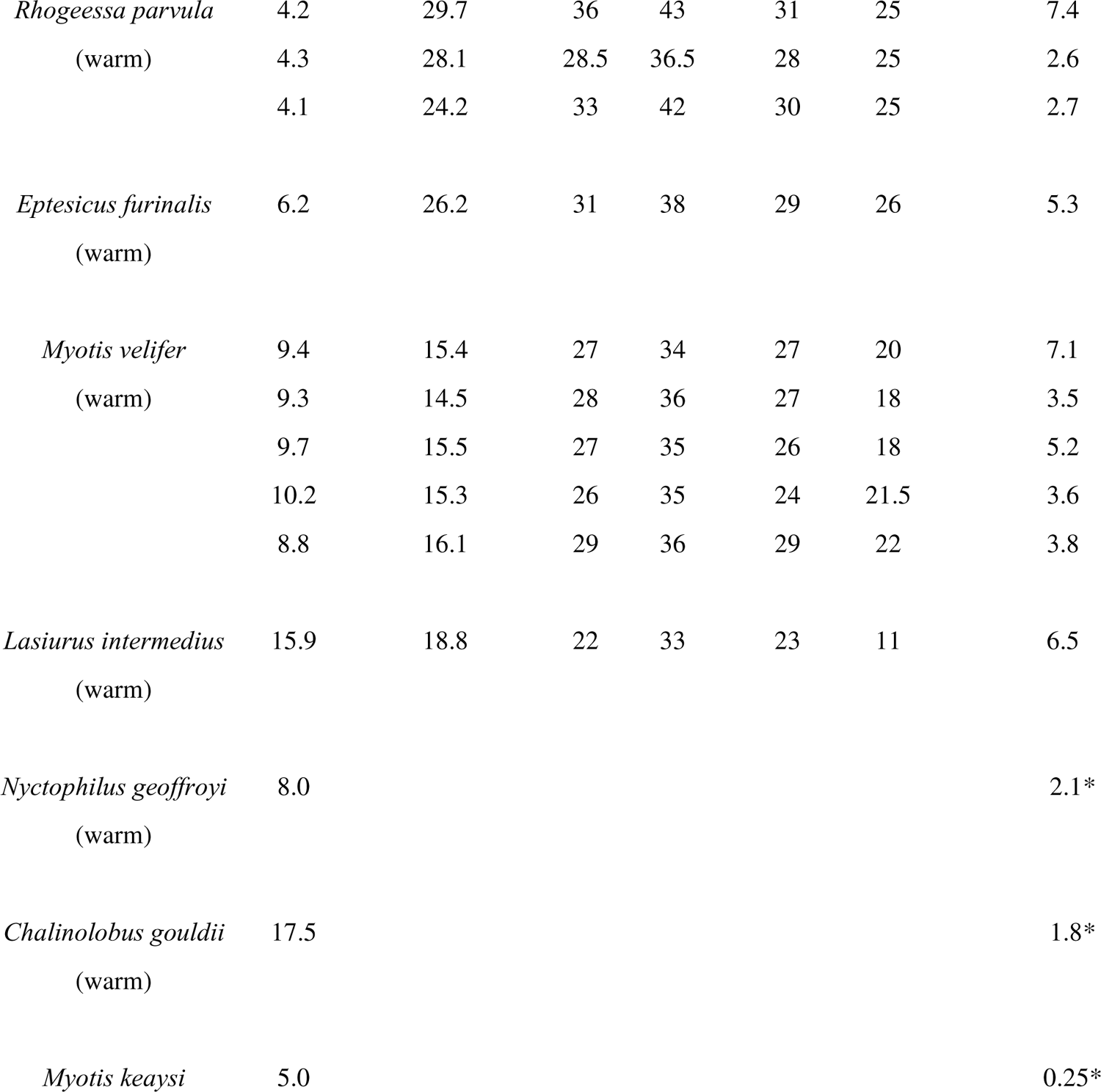
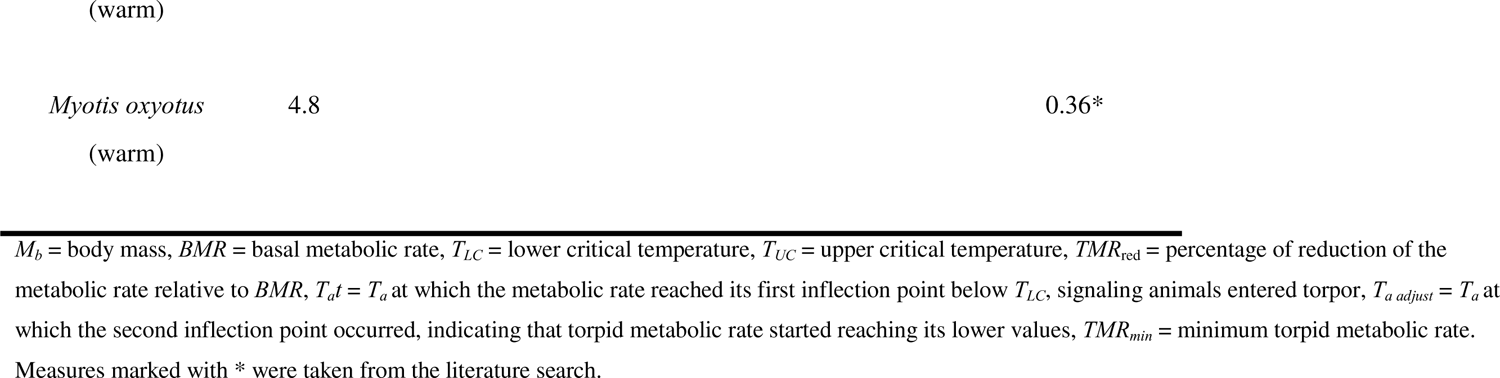
Thermal energetics of bats from the family Vespertilionidae living in cold and warm climates of central Mexico. Measurements were taken by using open flow respirometry of postabsorptive adult non-reproductive males during their resting phase from ∼ 10:00 to 19:30 hours.

Age was determined by observing the epiphyseal space of the fourth metacarpal bone of the third and fifth fingers, since juvenile individuals have some visible space. Reproductive condition of males was determined by observing the testes, which increase in size during spermatogenesis (Wilkinson and Brunet-Rossinni, 2009). Bats were captured under permission of the Wildlife Department granted to our institution (SEMARNAT: SGPA/DGVS/06795/21) and the ethics committee of the University of Tlaxcala.

Once captured, bats were placed in cloth bags and transferred to captive conditions near the study sites where they were placed in individual flight cages (75 cm x 75 cm x 75 cm) with 12/12 light/dark cycles, a *T_a_* of ∼28 °C (near the *TNZ* of bats), and a relative humidity above 50 %.

### Determination of thermal energetics in euthermic bats

Torpor is defined as a reduction of metabolic rate below *BMR* (Geiser and Ruf, 1995; Geiser, 2004), and *BMR* is measured between the *T_LC_*and *T_UC_* displayed by individuals (Withers et al., 2016). So, we initially measured *BMR*, *T_LC_*, and *T_UC_*of bats during normothermia by quantifying their resting metabolic rate (in mL O_2_ h^-1^) over a *T_a_* range of 8 to 43 °C. Measurements were taken ∼ 12 hours post-capture in a postabsorptive state during the resting phase (i.e., from ∼ 10:00 to 19:30 hours) (Genoud et al., 2018). Thermal traits were measured by indirect calorimetry, estimating O_2_ consumption and CO_2_ production using an open-flow respirometer (FoxBox®; Sable Systems International, Las Vegas). Measurements were done following Medina-Bello et al. (2023). To do this, each bat was placed in a 410 mL metabolic chamber inside a temperature-controlled cabinet (PELT5®; Sable Systems International, Las Vegas, USA). The chamber had plastic mesh on the walls and the ceiling to allow bats to hang upside down as in natural conditions, but it was too small for them to fly. This allowed us to minimize variation in the measurements due to the individuals’ movements. The temperature inside the cabinet was controlled to within ± 0.5 °C. Measurements were recorded when the readings reached an asymptote after the *T_a_* stabilized at ± 0.5 °C of each experimental *T_a_*. Flow rates (FR) of dry CO_2_ - free air scrubbed with Drierite (calcium sulfate for humidity) and Ascarite (sodium hydroxide for CO_2_) passed upstream with the air pushed through the respirometry chamber at a rate between 180- and 350-mL min^-1^ depending on the bats’ metabolic rate. Flow rates were estimated following Lighton and Halsey (2011) using the formula: FR= VO_2_/ ΔO_2_

Where VO_2_ corresponds to the predicted oxygen consumption of the experimental animal and ΔO_2_ is the difference in fractional concentration between the incurrent and excurrent O_2_. VO_2_ was estimated assuming that bats’ metabolic rates increase with *M_b_* with a slope of 0.744 (log_e_ BMR (mL O_2_ h^-1^) = 1.0895 + 0.744 log_e_ *M_b_* (g)) (Speakman and Thomas 2003). ΔO_2_ was taken from our FoxBox analyzer (0.0005 of ΔO_2_ expressed as fractional concentration). Excurrent air was dried with Drierite, and fractional concentrations (%/100) of both oxygen (F_e_0_2_) and CO_2_ (F_e_ C0_2_) were measured every second. We placed an empty hermetic chamber of the same size (410 mL) inside the *T_a_*cabinet to obtain reference measurements, since we did not have a system that allowed us to shift the airflow between the empty chamber and the animal chamber with the same respirometer. The empty chamber was connected to a second respirometer of the same brand (FoxBox®, Sable Systems International, Las Vegas, Nevada, USA) with the same type of sensors (O_2_ fuel cell and CO_2_ infrared sensors), and the same flow rate as that of the experimental chamber. Both respirometers were calibrated simultaneously and showed the same pattern of oxygen readings. Data from respirometers, including O_2_ and CO_2_ readings and data from the flowmeter and thermocouple, were sent to computers through Sable Systems-UI2 ports running Expedata software.

We placed each bat inside the metabolic chamber for 60 min at a *T_a_* of 28 °C before taking any measurement to allow each bat to recover from the stress of handling. The first *T_a_* tested was 28 °C, which we have previously determined falls within the *TNZ* of all of the bat species studied here (Medina-Bello et al., unpublished data). We measured the metabolic rate of individuals for 30 min at 28 °C, then decreased the *T_a_* to 8 °C, taking measurements for 5 min for each drop in 1 °C and 30 min for each 5 °C drop (i.e., at 23, 18, 13, and 8 °C). Subsequently, we increased the chamber’s *T_a_* to 28 °C for 30 min before raising it to 43 °C. Measurements were recorded for 5 min for each 1 °C increase and 30 min for each 5 °C increase (i.e., at 33, 38, and 43 °C). It took 5 to 10 min for the *T_a_* to stabilize for each change in *T_a_* within the metabolic chamber. The metabolic measurements we obtained from the animals at each 1 °C increment allowed us to obtain a more precise estimation (1 °C rather than a 5 °C accuracy) of the *BMR*, *T_LC_*, and *T_UC_*. One individual was measured in the set of *T_a_*’s at each time. We increased the *T_a_* inside the metabolic chamber and repeated the measurements if the bats reduced their metabolic rate below their *BMR* and intended to use torpor. With this methodology, bats from all the sites increased their resting metabolic rate when the *T_a_*fell below their *TNZ*, indicating that they defended normothermia. At the end of these procedures, bats were provided with mealworms (*Tenebrio molitor*) and water *ad libitum* and kept in captivity for the torpor experiments.

### Determination of thermal energetics during torpor

To induce bats to use torpor, we deprived the individuals of food for 36 h before conducting any experiment (Ruf et al., 1993). This period of food deprivation mirrors the short-term food restrictions that bats may encounter in the field (Geiser, 2021). With this methodology, all bats used torpor during our trials. During captivity, bats were kept in the previously described flight cages. After the 36 hrs., we measured the metabolic rate of individuals during their resting phase (from 8:00 to 17:00 hours). We placed each bat in the metabolic chamber for 60 min at a *T_a_* of 28 °C to allow them to recover from handling. As we were particularly interested in identifying the inflection points below the *BMR* that defined torpor traits, the first *T_a_* we tested was the mean *T_UC_* we calculated for each species. This allowed us to maintain the bats in their thermoneutral zone before they reached their *T_LC_*and used torpor. One individual was measured at a time. We measured the metabolic rate of individual for 75 min, then we decreased the *T_a_*to 8 °C, with measurements taken for 7 min for each 1 °C drop and 75 min for each 5 °C drop. Using this protocol, we were able to measure torpid metabolic rate for five to six hours continuously, which allowed us to obtain precise measurements of torpor traits in our study species (Geiser, 2021). Since individuals exhibited a metabolic rate below the *BMR* during torpor, we adjusted the FR between 110 mL min^-1^ and 180 mL min^-1^ to achieve accurate measurements of *T_a_t*, *T_a adjust_*, and *TMR_min_*. In both euthermic and torpor measurements, we obtained the *M_b_* by weighing the individuals at the beginning of the experiments by using the electronic balance previously described. These measurements were used for data analyses. No bats died during the experiments. The individuals were fed, provided with water, and released at their captured sites the night following the conclusion of the experiments.

### Data analyses Metabolic rate

All data analyses were conducted using RStudio software (version 1.2.5042). We measured the metabolic rate of bats through oxygen consumption (VO_2_) (in mL O_2_ h^-1^) for each bat tested at each *T_a_*. The metabolic rate was calculated using the formula proposed by Lighton (2018):

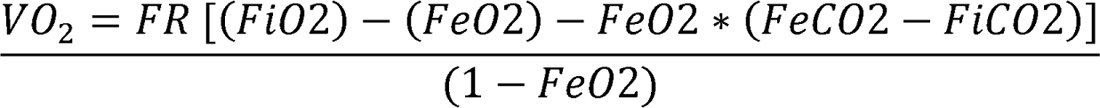

Where FR is the flow rate (in mL min^-1^), FiO_2_ is the fractional concentration of O_2_ in the incurrent (baseline) air, FeO_2_ is the fractional concentration of O_2_ in the excurrent air, FiCO_2_ is the fractional concentration of CO_2_ in the incurrent air, and FeCO_2_ is the fractional concentration of CO_2_ in the excurrent air. To conduct our analyses, we used the mean values of the metabolic rate obtained from the measurements of the experimental *T_a_*’s and the mean values obtained from the animals at each °C. We defined *BMR* (in mL O_2_ h^-1^) as the mean metabolic rate of the bats between *T_LC_* (°C) and *T_UC_* (°C) (Genoud et al., 2018, Geiser, 2021). Critical temperatures were obtained using two-phase regression (TPR) with the ‘chngptm’ function (‘chngptm’ package). TPR identifies thresholds of abrupt changes in the relationship between the dependent and independent variables (Nickerson et al., 1989). In bats, TPR has been used to estimate thermal traits including *BMR*, *T_LC_*, and *T_UC_* (see Willis et al., 2005a and 2005b and Machado and Soriano, 2007; among others). In our models, the metabolic rate of the bats was the dependent variable, and *T_a_* the independent one.

To determine the torpor traits of bats, we first fitted a simple linear regression model using the ‘lm’ function of the ‘stats’ package. In this model, the response variable was the mean value of the metabolic rate of bats at different *T_a_* below the *T_UC_*, and *T_a_* was the explanatory variable. We calculated the inflection points of the regression model by using the ‘davies.test’ function of the ‘segmented’ package (Muggeo, 2008). In this calculation, the first inflection point indicated *T_a_t* of bats, signaling that they went into torpor. The second inflection point indicated *T_a adjust_*. Finally, we obtained the minimum torpid metabolic rate (*TMR_min_*) by calculating the mean of the torpid metabolic rate for ∼ 2 hrs. below *T_a adjust_*. Because some authors have measured *TMR_min_* for certain bat species of the family Vespertilionidae, we conducted a systematic search in Google scholar and the Web of Knowledge to look for those values to include them in our data. The search was done using the keywords: “torpor Vespertilionidae”, “vespertilionid minimum metabolic rate”, “torpor bats Vespertilionidae”. We found calculations of *TMR_min_* for 14 species of bats.

However, only data from *Nyctophilus geoffroyi*, and *Chalinolobus gouldii* (Hosken and Whiters, 1999), *Myotis myotis* (Hanus, 1959), *Myotis keaysi*, and *Myotis oxyotus* (Manchado and Soriano, 2007) were useful for our analysis, since those were the only studies in which *TMR_min_* was measured in short-term experiments that were comparable to our study design.

### Effect of M_b_ and climate on the torpid metabolic rate of bats

We evaluated the relationship of *M_b_* and climate with 1) *BMR*, 2) percent reduction of the metabolic rate below *BMR* (*TMR*_red_), and 3) torpor traits of bats. To conduct our analyses, we used phylogenetic mixed-effect models, using the ‘gls’ function of the ‘nlme’ package’. We employed a Brownian correlation structure based on the phylogenetic tree published by Amador et al. (2018) for the order Chiroptera. This correlation accounts for the shared evolutionary history among the bat species (Paradis and Schliep, 2019). We used the ‘drop.tip’ function of the ‘ape’ package to trim the tree to the 590 species of bats of the family Vespertilionidae and two species of the family Molossidae (*Eumops floridanus* and *Eumops glaucinus*) that were used as outgroups. To meet model assumptions of normality, we transformed the *BMR*, *T_a_t* and *T_a adjust_* data using the ‘bestNormalize’ function of the ‘bestNormalize’ package. Given that phylogenetic models have been reported to offer a better fit than conventional linear regressions (Riek and Geiser, 2013), we created simple linear regressions and compared them with our phylogenetic models using the ‘lrtest’ function from the ‘lmtest’ package. In the models, *BMR*, *TMR*_red_, and torpor energetics (i.e., *T_a_t*, *T_a adjust_* and *TMR_min_*) were the dependent variables, while *M_b_* and climate (classified as categorical—either cold or warm) were the independent ones. To conduct our analyses, we utilized the *M_b_*, *TMR*_red_, *T_a_t*, *T_a adjust_* and *TMR_min_*measurements we obtained from our experiments. For the *TMR_min_*, we also integrated the data we gathered from our bibliographic search. The slopes of the relationships between explanatory and response variables (β_0_) along with their respective standard error were extracted from model summaries. As our primary interest was to assess the effect of the combination of *M_b_*and climate on the torpor energetics of bats, we tested the interaction between these explanatory variables. Notably, we found significant interactions between *M_b_* and climate in all our models. Given that these interactions provided a more comprehensive representation of the relationship we intended to describe, we present the results in terms of the interactions.

For the models, we constructed a vector containing the position of each measurement of the corresponding bat species in the phylogenetic tree using the ‘match’ function from the ‘base’ package. We employed this vector as a grouping factor and to account for the heteroscedasticity of the repeated measures obtained from our measurements using the ‘weight’ and ‘varIndent’ functions from the ‘sjstats’ and the ‘nlme’ packages, respectively. As *P* values for parameter estimates within mixed models are not as straightforward as those obtained from linear regressions, we obtained the *P* values from our phylogenetic models by comparing the likelihood ratio against null models using the ‘lrtest’ function from the ‘lmtest’ package. Tests were considered significant at an Iii value ≤ 0.05.

## Results

In this section, data are presented as the mean ± standard error unless otherwise noted. We obtained euthermic and torpor traits from 33 individuals belonging to 11 bat species from the family Vespertilionidae and *TMR_min_* for five species from the bibliography search (Table 2). Bats from the warm climates were in general smaller (7.05 ± 1.03 g) and presented higher *BMR* (21.01 ± 1.63 mL O_2_ h^-1^), *TMR*_red_ (76.78 ± 2.82 %), *T_a_t* (27.89 ± 1.19 °C) and *T_a adjust_*(22.50 ± 1.19 °C) than bats from the cold climates (11.87 ± 1.42 g, 16.58 ± 0.86 mL O_2_ h^-1^, 67.98 ± 2.01 %, 27.89 ± 1.19 °C, 22.50 ± 1.19 °C, for the *M_b_*, *BMR*, *TMR*_red_, *T_a_t*, and *T_a adjust_*, respectively). Nevertheless, bats from the warm climates presented lower values of *TMR_min_* (3.72 ± 0.48 mL O_2_ h^-1^) than bats from the cold climates (5.02 ± 0.41 mL O_2_ h^-1^).

All phylogenetic models fit the data better than conventional ones (Table 3). Therefore, we utilized the phylogenetic models to interpret our results (Freckleton 2009). Our results are conceptually summarized in Fig. 3. In our models we found a positive relationship between the interaction of *M_b_*and climate with the *BMR* of bats (β_0_ = 0.10 ± 0.01 and β_0_ = 0.09 ± 0.00 for bats from the warm and cold climate, respectively) (*X^2^* = 6.05, df = 0, p < 0.0001), such that *BMR* increased more strongly with increasing *M_b_*in bats from warm climates than those from cold climates (Fig. 4A). We also found a remarkably strong relationship between *T_LC_* and *T_a_t* of bats (r^2^ = 0.91, t = 27.1, df = 31, p < 0.0001) (Fig. 4B). When individuals entered torpor, they experienced *TMR*_red_ between 50 % and 90 % below *BMR*. This reduction was negatively affected by the interaction of *M_b_* with climate (β_0_ = −0.52 ± 0.04 and β_0_ = −0.45 ± 0.15 for bats from the warm and cold climate, respectively) (*X^2^* = 8.89, df = 0, p < 0.0001) (Fig. 4C), such that *TMR_red_* decreased with increasing *M_b_* more strongly in bats from warm climates than in those from cold climates. We also found a negative relationship between the interaction of *M_b_* with climate with *T_a_t* (β_0_ = −0.09 ± 0.00 and β_0_ = −0.19 ± 0.01 for the warm and the cold climate, respectively) (*X^2^* = 4.58, df = 0, p < 0.0001) (Fig. 4D), as well as the *T_a adjust_* (β_0_ = −0.03 ± 0.01 and β_0_ = −0.10 ± 0.01 for the warm and the cold climate, respectively) (*X^2^* = 29.8, df = 0, p < 0.0001) (Fig. 4E). Thus, both *T_a_t* and *T_a adjust_* decreased more strongly with increasing *M_b_* in bats from cold climates than in those from warm climates. Finally, we found a positive relationship between the interaction of *M_b_* with climate with the *TMR_min_* of bats (β_0_ = 0.19 ± 0.02 and β_0_ = 0.17 ± 0.02, for bats from the warm and cold climate, respectively) (*X^2^* = 3.34, df = 0, p < 0.0001) (Fig. 4F), such that *TMR_min_* increased with increasing *M_b_* more strongly in bats from warm climates than in those from cold climates.

**Figure 3.**
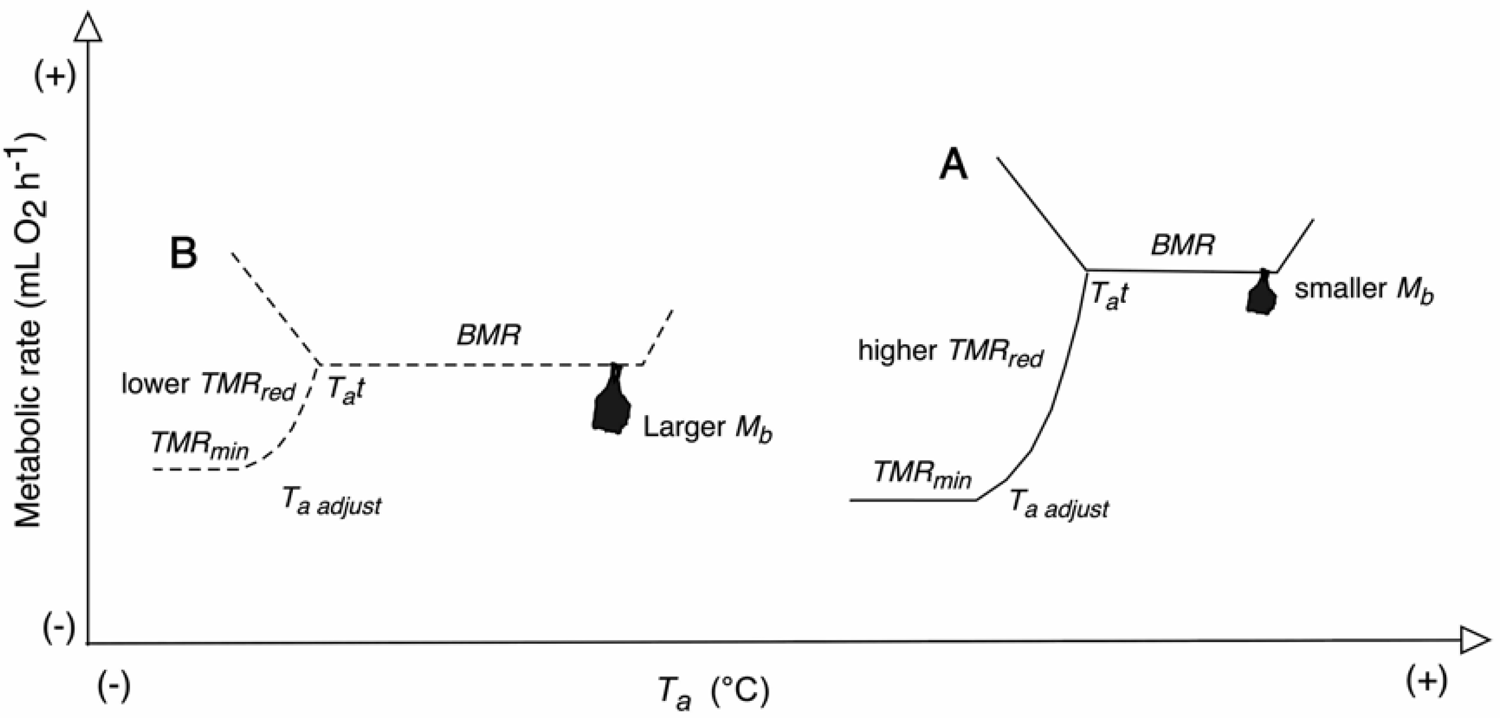
Summary of the thermal energetics of bats from warm (A) and cold (B) climates of central Mexico obtained in this study. Here we show how bats from the warm climates were smaller and showed higher basal metabolic rate (*BMR*) and torpid metabolic rate reductions (*TMR_red_*), warmer temperatures at which bats entered torpor (*T_a_t*) and reached its lower values (*T_a adjust_*) compared to bats from the cold climates. Nevertheless, because bats from the warm climates presented higher *TMR_red_*, minimum torpid metabolic rates (*TMR_min_*) were lower than bats from the colder ones.

**Figure 4.**
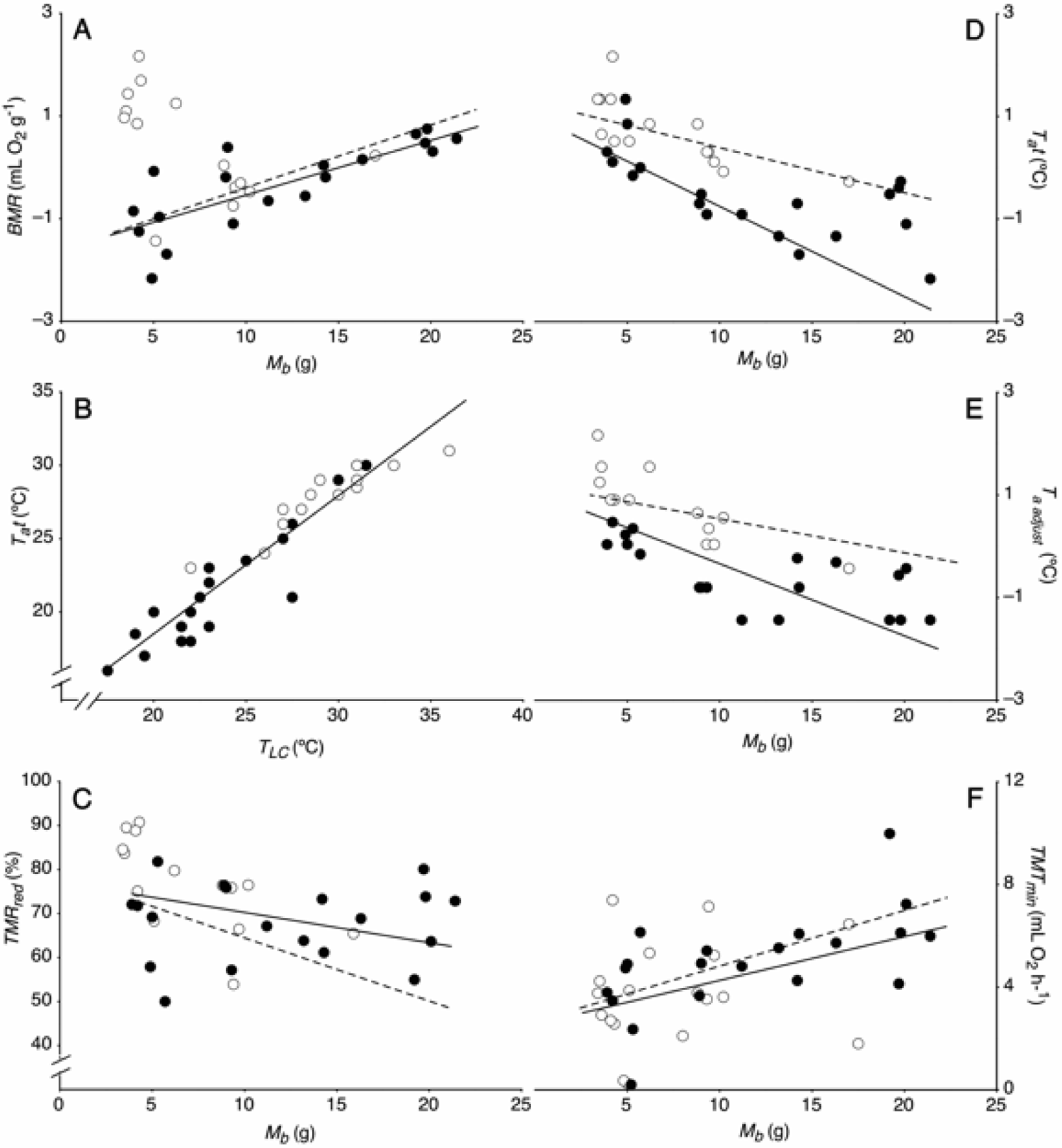
Effect of body mass (*M_b_*) and climate (either warm-open symbols- or cold-closed symbols-) on basal metabolic rate (*BMR*), lower critical temperature (*T_LC_*) during euthermia, and torpor energetics (*T_a_* at which animals entered torpor (*T_a_t*), and the inflection point (*T_a adjust_*) where torpid metabolic rate started reaching its lower values (*TMR_min_*)) from vespertilionid bats living in central Mexico. Regression lines (dashed lines-warm climates-, and continuous lines-cold climates-) were fitted by using the estimated parameters from the phylogenetic models.

**Table 3.** Statistical comparison between models used to evaluate the relationship between *M_b_* and climate in torpor traits of bats of the family Vespertilionidae. Comparisons were made by using the ‘lrtest’ function from the ‘lmtest’ package in RStudio (version 1.2.5042).

| Model structure | Model type | AIC | Statistics |
| --- | --- | --- | --- |
| $BMR \sim M_b : \text{climate}$ | phylogenetic | - 426.5 | $X^2 = 658.4$, $df = 10$, $p < 0.0001$ |
|  | conventional | 211.8 |  |
| $TMR_{\text{red}} \sim M_b : \text{climate}$ | phylogenetic | - 268.1 | $X^2 = 2243.4$, $df = 10$, $p < 0.0001$ |
|  | convectional | 248.7 |  |
| $T_{at} \sim M_b : \text{climate}$ | phylogenetic | - 2152.9 | $X^2 = 2250.0$, $df = 10$, $p < 0.0001$ |
|  | conventional | 77.0 |  |
| $T_{a \text{ adjust}} \sim M_b : \text{climate}$ | phylogenetic | -2142.7 | $X^2 = 2241.2$, $df = 10$, $p < 0.0001$ |
|  | conventional | 78.4 |  |
| $TMR_{\text{min}} \sim M_b : \text{climate}$ | phylogenetic | -2053.1 | $X^2 = 2255.8$, $df = 16$, $p < 0.0001$ |
|  | conventional | 170.7 |  |

$M_b$ = body mass (g), $BMR$ = basal metabolic rate ( $\text{mL O}_2 \text{ h}^{-1}$ ); $TMR_{\text{red}}$ = percentage of reduction of the metabolic rate relative to $BMR$ (%), $T_{at} = T_a$ at which the metabolic rate reached its first inflection point below $T_{LC}$ , signaling animals entered torpor ( $^{\circ}\text{C}$ ), $T_{a \text{ adjust}} = T_a$ at which the second inflection point occurred, indicating that torpid metabolic rate started reaching its lower values ( $^{\circ}\text{C}$ ), $TMR_{\text{min}}$ = minimum torpid metabolic rate ( $\text{mL O}_2 \text{ h}^{-1}$ )

## Discussion

In this study, we found that phylogenetic models were a better fit for our data than conventional linear regressions. This finding is consistent with previous research on thermal energetics during euthermia and torpor for some species of birds and mammals, including bats (e.g., Rezende et al., 2004; Cory Toussaint and McKechnie, 2012; Geiser, 2013; Riek and Wolf, 2020). Because phylogenetic models account for shared evolutionary relationships among taxa (Paradis and Schliep, 2019), this makes them a more realistic tool for interpreting the relationships among morphological, ecological, and physiological traits (Garland et al., 2005; Freckleton, 2009). Therefore, their use in comparative studies should be promoted.

In our models, we found a positive relationship between the interaction of *M_b_* and climate with the *BMR* of bats. This can be explained by the combination of several factors associated with the morphology and physiology of individuals. First, we found smaller bats in warm climates compared to the colder ones. A similar trend in *M_b_* variation has been observed among bats inhabiting sites with a variety of *T_a_*’s, suggesting adaptations in *M_b_* linked to thermoregulation. For example, Alston et al. (2023) compiled data from ∼ 31,000 individuals across 20 bat species captured over a decade in North America. The authors found that bats from colder climates typically exhibited larger *M_b_* than those from warmer environments. In colder climates, larger *M_b_* can be advantageous because the surface-area-to-volume ratio decreases with increasing *M_b_*, which may help individuals to lose less metabolic heat to the environment compared to their smaller counterparts (Austad and Fischer, 1991; Speakman, 2005). Second, smaller individuals exhibited lower *BMR* values than larger ones. This result aligns with findings in other mammals and birds worldwide (Hayssen and Lacy, 1985; Symonds and Elgar, 2002; White and Seymour, 2003; White et al., 2007, 2009; Packard and Birchard, 2008; among others) and can be explained simply because larger animals possess more metabolizable tissue than the smaller ones (Williams and Tieleman, 2000). However, bats from the warm climates had higher *BMR* values than those from the cold climates. Because smaller animals require greater metabolic heat production to maintain a high *T_b_* due to their higher surface-area-to-volume ratio, this could lead to elevated *BMR*’s in bats from the warmer sites (Speakman and Thomas, 2003; Clarke et al., 2010). Third, individuals from the warm climates increased more strongly their *BMR* with *M_b_* than those from the cold climates. In our trials, we found exceptionally high *BMR* values in very small *M_b_* species like *Rhogeessa parvula*, *R. alleni* and *E. furinalis* from the warmer sites, which presented *BMR* values four to six times higher than what would be expected for their *M_b_* and as much as double the *BMR* of *Myotis californicus* and *M. volans*, two bat species of similar *M_b_* from the colder sites. This could be the reason because we found a higher slope in the relationship between *M_b_*and climate with the *BMR* of bats for the warm climates. Differences in *BMR* have been associated to organ size in birds and mammals (Kersten and Piersma, 1987; Konarzewski and Diamond, 1995; Nespolo et al., 2002). In bats, there is scant information about this topic, yet larger brains have been correlated to higher *BMR*’s (McNab and Köhle, 2017). It has been also demonstrated that bats living in more complex environments tend to present larger brains than those living in simpler environments (Safi et al., 2005). Because warmer sites tend to present more vegetation compared to colder sites (and hence more complexity) (Gaston, 2000; Kraft and Ackerly, 2014), bats from warmer sites should have presented larger brains, leading to higher metabolic rates. Although differences in brain mass can explain the differences we observed here, this theme calls for further exploration. Higher *BMR*’s would be more sustainable in warmer climates where prey abundance tends to be higher, and *T_a_* is typically more stable than in colder environments (Wolda, 1978). Nonetheless, the use of torpor might help individuals in both warm and cold climates achieve energy balance when facing energy constraints due to both *T_a_* and food availability.

In our measurements, we found a strong correlation between *T_LC_*’s and *T_a_t* experienced by bats. This suggests that our study species efficiently conserved energy when faced with energy constraints due to the 36-hour food deprivation before experiments, combined with exposure to *T_a_*’s below their *T_LC_*. Lower critical temperature represents the *T_a_* at which individuals start expending energy to generate heat and maintain a stable *T_b_* that is in most cases higher than *T_a_*(Fristoe et al., 2015). However, under energy-restricted conditions, minimizing metabolic expenditures becomes crucial (Vuarin et al., 2014, 2015). To achieve energy balance, our study species employed torpor, significantly reducing their metabolic rate to 50 % to 90 % below their *BMR* (i.e., *TMR_red_*). Similarly, other daily heterotherms from the orders Rodentia, Insectivora, Chiroptera and Carnivora have been observed displaying a *TMR_red_* ranging from 43 % to 96 % when using torpor in both laboratory and natural conditions (Ellison, 1992; Holloway and Geiser, 1995; Geiser, 2004 and the references cited there; Geiser et al., 2019). Nevertheless, in our experiments we found that *TMR_red_* was negatively affected by the interaction of *M_b_* and climate. In our trials, we found high *TMR_red_*values in smaller *M_b_* bats compared with their larger counterparts. Geiser (2021) has proposed that higher *TMR_red_* in small animals has likely evolved to maximize energy-savings. Differences in *TMR_red_*can be attributed to the evaporative cooling capacity experienced by animals when entering torpor, which could be influenced by their *M_b_*. Smaller animals tend to exhibit elevated resting metabolic rates (*RMR*) at low *T_a_*. When these individuals enter torpor, their *RMR* undergoes substantial reduction towards torpid values. This reduction, combined with the large surface-area-to-volume ratio, leads to a rapid decrease in the *T_b_*of individuals, promoting a steep and deep decline in metabolism. This could be the reason because *R. parvula*, *R. alleni* and *E. furinalis*, which presented the highest *BMR* values and smallest *M_b_*among our study species exhibited the highest *TMR_red_* values (averaging from 79 to 85 %) in our experimental trials. This can also explain why the slope of the relationship between *TMR_red_* and the interaction of *M_b_* and climate was higher for bats from the warmer climates compared to the colder ones. However, in our experimental trials, all bats were confronted with *T_a_*’s below their *T_LC_* when animals were within their *TNZ* (see methods section).

Under these circumstances, physiological inhibition coupled with the abandonment of *T_b_* regulation could have contributed to significant reductions in the metabolic rate of bats, especially of those of the warm climates (Reher and Dausmann, 2021). While physiological inhibition has been reported more for hibernating animals rather than daily heterotherms (Geiser, 1988; Lyman, 1982), it is plausible that a similar phenomenon has occurred in our study species. Nevertheless, this topic needs further exploration. Larger bats, which are adapted to colder climates, exhibit lower *T_LC_* values (Speakman and Thomas, 2003; Geiser, 2021), and the disparity between *RMR* and *BMR* tends to be less pronounced. In our study sites, bats from colder climates exhibited a mean *T_LC_* of 23.7 ± 0.89 °C compared to the 29.3 ± 0.90 °C observed in bats from warmer climates. As larger bats enter torpor, their metabolic rate drops may be less pronounced. Due to this condition and their lower surface-to-volume ratio, this leads to slower cooling rates that could result in higher torpid metabolic rates compared to bats from warmer sites (Geiser, 2004). This also can explain the lower slope of bats from the colder climates we found for the relationship between *TMR_red_* and the interaction of *M_b_* and climate compared to the warmer ones. Cooling rates have important co-evolutionary implications in roost selection by bats. Czenze et al. (2021) demonstrated that bat species occupying poorly buffered roosting sites exhibited higher evaporative cooling capacity compared to those using well-isolated ones. This can explain why small *M_b_* bats such as *R. parvula*, *R. alleni*, and *E. furinalis* roost in poor isolated roosts, such as underneath exfoliating bark and inside shallow tree cavities in our study sites (Mies et al., 1996; Roots and Baker, 2007). Because roosting places are closely related to torpor use in bats (Alston et al., 2022), this topic needs more investigation.

In this study we identified a negative relationship between the interaction of *M_b_* and climate with the *T_a_t* and *T_a adjust_*of bats, along with a positive relationship with *TMR_min_*. In our experiments, smaller *M_b_* bats presented higher values of *T_a_t* and *T_a adjust_* and lower values of *TMR_min_* compared to their larger counterparts. This can explain why bats from warm climates in nature are able to use torpor at high *T_a_*’s in the tropics (e.g., Turbill et al., 2003; Geiser et al., 2011; Liu and Karasov, 2011). These results can be explained by the strong correlation we found between *T_LC_* and *T_a_t* of bats, as well as differences in *TMR_red_* observed among bats of the different climates. As *T_LC_* inversely relates to *M_b_*, smaller *M_b_* bats display higher *T_LC_* values leading to elevated *T_a_t* compared to larger ones. Furthermore, due to the sequence of torpor events, *T_a adjust_* occurred as the second inflection point after *T_a_t*, resulting in warmer *T_a adjust_* values in smaller bats as well. Interestingly, we observed that smaller bats presented higher *TMR_red_*values, which ultimately contributed to lower *TMR_min_* than their larger counterparts. Given that bats from warmer sites had lower *M_b_*, this may account for the differences we found in bats from cold and warm climates in our study sites.

## Conclusion

This investigation underscores the significant role of the interaction between *M_b_* and climate on the torpor traits of bats. However, more comprehensive research on this subject needs to be conducted. While numerous avenues for investigation exist, we propose several ideas related to the results we found. For instance, investigations should attempt to understand why extremely small bats endure significantly cold climates over extended periods of time. For example, *Myotis californicus* and *M. melanorhinus* (∼ 4 g), and *M. volans* (∼ 6 g) are found in La Malinche National Park for a significant portion of the year (Ayala-Berdon et al., 2017). Despite limited reports of hibernation (Aguilar-Rodríguez et al., 2021) the utilization of torpor by these species remains largely unexplored. Further investigation should also explore regional disparities in torpor usage among bats with extensive geographic ranges. In this regard, *Eptesicus fuscus* and *Lasiurus borealis* from North America presented lower *T_a_t*’s in higher latitudes where *T_a_* tends to be colder than lower latitudes where *T_a_* is typically warmer (Dunbar and Brigham, 2010). Although these differences have not been assessed in different climates, *M. velifer* presents higher *BMR*, thermal conductance, and *T_LC_*and *T_UC_* during euthermia when captured from warm sites compared to a cold site in central Mexico (Medina-Bello et al., 2023). Nevertheless, no information from torpor traits from this species has been collected. Therefore, more work in this area needs to be done.

## Acknowledgments

This work supported by the program CONACYT FOSEC CB2017-2018 (A1-S-39572) granted to JAB. We are grateful to L. Orozco-Lugo for her help capturing bats in Quilamula, to R. Vázquez-Fuerte for her comments on the manuscript and the ejido of Moxolahuac and the Quilamula Biological Station for logistical support.

## Competing interests

The authors declare there are no competing interests

## Funding

This investigation was supported by the program CONACYT FOSEC CB2017-2018 (A1-S-39572) granted to JAB. Supporters had no participation in study design, data collection, analysis, or the writing of the manuscript.

## Data availability

All data obtained in this work is fully presented in the manuscript.

## Notes

### Competing Interest Statement

The authors have declared no competing interest.

